# Development of Reverse Transcription Loop-mediated Isothermal Amplification (RT-LAMP) Assays Targeting SARS-CoV-2

**DOI:** 10.1101/2020.03.09.983064

**Authors:** Gun-Soo Park, Keunbon Ku, Seung-Hwa Baek, Seong Jun Kim, Seung Il Kim, Bum-Tae Kim, Jin-Soo Maeng

## Abstract

Epidemics of Coronavirus Disease 2019 (COVID-19) now have more than 100,000 confirmed cases worldwide. Diagnosis of COVID-19 is currently performed by RT-qPCR methods, but the capacity of RT-qPCR methods is limited by its requirement of high-level facilities and instruments. Here, we developed and evaluated RT-LAMP assays to detect genomic RNA of SARS-CoV-2, the causative virus of COVID-19. RT-LAMP assays in this study can detect as low as 100 copies of SARS-CoV-2 RNA. Cross-reactivity of RT-LAMP assays to other human Coronaviruses was not observed. We also adapted a colorimetric detection method for our RT-LAMP assay so that the tests potentially performed in higher throughput.

## Introduction

SARS-CoV-2 is the causative viral pathogen of COVID-19 of which the outbreaks resulted in 106,893 confirmed cases involving 3,639 deaths over 103 countries, areas or territories as of 07 March 2020.[1] As the name suggests, SARS-CoV-2 is closely related to a group of severe acute respiratory syndrome-related coronaviruses (SARSr-CoV), namely subgenus Sarbecovirus, showing 96% identity to a bat coronavirus [2, 3].

Diagnosis of COVID-19 can be done through CT scan of suspicious patients and confirmatory lab test is performed using RT-qPCR methods [4-8]. Although RT-qPCR methods are used as the gold-standard for detection of pathogens due to its high sensitivity and specificity, it still have some caveats. Briefly, lab-level facility such as reliable supply of electricity, expensive instruments and trained personnel are required to properly perform RT-qPCR tests. These restrictions hinder to utilize RT-qPCR methods for various point-of-care where pathogen detection might be required [9].

To overcome such cost-restriction of RT-qPCR and still detect pathogens’ nucleic acids, isothermal amplification methods have been developed [10]. Among such methods, Loop-mediated isothermal amplification (LAMP) method has some advantages to be applied for point-of-care test (POCT) [11]. Well optimized LAMP assay shows sensitivity comparable to that of PCR, less than 10 copies per reaction [12]. Intercalating fluorescent dyes are compatible with LAMP reaction so that we can observe amplification in real-time [13]. Because amplification efficiency of LAMP reaction is very high, changes in reaction mixture components made it possible to detect the result with colorimetric detection methods [14-16]. Moreover, unpurified sample can be directly used for LAMP [17-19]. This indicates that high-throughput test is possible when use of unpurified specimen is combined with non-instrumental (*e.g*. colorimetric) detection.

In this study, we designed and evaluated one-step reverse transcription LAMP (RT-LAMP) methods to detect SARS-CoV-2. We provide a pair of LAMP primer sets that is specific to SARS-CoV-2 and accompanying optimized reaction conditions. To achieve colorimetric detection of LAMP reaction, leuco crystal violet (LCV) method is applied [15].

## Materials and Methods

### Viral RNA preparation

SARS-CoV-2 viral RNA was prepared as previously described [20]. hCoV-229E and hCoV-OC43 viral RNA were isolated from culture media of infected MRC-5 cells (ATCC^®^ CCL-171™). MERS-CoV RNA was isolated from cell pellet lysate of infected Vero cells (ATCC^®^ CCL-81™).

### Viral RNA titration

*In vitro* transcribed standard RNA for SARS-CoV-2 was prepared as previously described [20]. To prepare standard RNA for hCoV-229E, first amplicon of PCR (Forward primer: 5’ – GCTAGTGGATGATCATGCTTTG – 3’, Reverse primer: 5’ – TGGGGCCATAAACTGTTCTATTAC – 3’) was cloned to pBluescript II KS (+) plasmid with *BamH*I and *Xho*I. Then, *in vitro* transcription template was prepared by restriction enzyme cut with *Bgl*I and *Xho*I and purification after agarose gel electrophoresis. For hCoV-OC43 and MERS-CoV, amplicons of PCR (OC43 forward primer: 5’ – AGCAACCAGGCTGATGTCAATACC – 3’, OC43 reverse primer: 5’ – AGCAGACCTTCCTGAGCCTTCAAT – 3’, MERS-CoV: UpE region [21]) were synthesized and cloned to pBIC-A plasmid (Bioneer). *In vitro* transcription template for hCoV-OC43 and MERS-CoV were prepared by restriction enzyme cut with *BamH*I-*Xho*I or *Ssp*I-*Xho*I, respectively. *In vitro* transcriptions were done with EZ™ T7 High Yield In Vitro Transcription kit (Enzynomics) as manufacturer’s instructions. RNA products were then purified using Agencourt RNAClean XP (Beckman Coulter). Standard RNA copy numbers were calculated from concentration measured by NanoDrop Lite (Thermo Scientific). All restriction enzymes were purchased from Enzynomics.

To evaluate genomic copy number of viral RNAs, dilutions of standard RNAs and viral RNAs in TE buffer (10 mM Tris-Cl, pH 8.0, 1 mM EDTA) are subjected to one-step RT-qPCR. RT-qPCR reactions were carried out using LightCycler 96 instrument and following reagents: Luna Universal One-Step RT-qPCR Kit (New England Biolabs, NEB) for hCoV-229E and hCoV-OC43, THUNDERBIRD Probe qPCR Master Mix (Toyobo) for MERS-CoV and Luna Universal Probe One-Step RT-qPCR Kit (NEB) for SARS-CoV-2.

### Reverse transcription

cDNA of SARS-CoV-2 was made using SuperScript IV Reverse Transcriptase (Invitrogen) following manufacturer’s instructions with modifications. Briefly, 10 pmol of random hexamer was used as reverse transcription primer and reaction was performed as follow: 20 minutes at 25°C, 30 minutes at 55°C and 10 minutes at 80°C.

### LAMP and RT-LAMP reaction

LAMP reaction was performed with reaction mixture containing following components: 1.6 μM FIP/BIP primers, 0.2 μM F3/B3 primers, 0.4 μM LF/LB primers, 1x Isothermal Amplification Buffer II (NEB, 20 mM Tris-HCl pH 8.8, 10 mM (NH_4_)_2_SO_4_, 150 mM KCl, 2 mM MgSO_4_, 0.1% Tween^®^ 20), 6 mM MgSO_4_ (NEB, final 8mM Mg^2+^), 1.4 mM each dNTP (Enzynomics), 0.4 μM SYTO-9 (Invitrogen) and 6 U Bst3.0 DNA polymerase (NEB) in total 15 μl reaction volume. For RT-LAMP, 10 U of SuperScript IV Reverse Transcriptase (Invitrogen) was added. For end-point colorimetric detection of LAMP reaction, tweaked version of 5x stock LCV solution[15] containing 0.5 mM Crystal Violet (Sigma, C0775), 60 mM Sodium Sulfite (Sigma, S0505) and 5 mM β-Cyclodextrin (Sigma, C4767) was directly added to LAMP reaction mixture to 1x concentration. When using WarmStart Colorimetric LAMP 2X Master Mix (NEB), same concentrations of primers and 0.4 μM SYTO-9 were added. Isothermal incubation and fluorescence signal measurement was performed using LightCycler 96 instrument (Roche) at 69°C with additional heat inactivation (5 minutes at 95°C) and melting curve analysis steps. Fluorescence signals were measured for every minute during 60 (screening and optimization) or 30 (Limit of Detection and cross-reactivity) minutes of incubation. Any changed conditions are specified for each experiment.

## Results

### LAMP Primer Design

Five SARS-CoV-2 sequences (MN908947, MN938384, MN988713, MN985325, and MN975262) and seven SARS-CoV sequences (NC_004718, AY613947, AY313906, AY559094, AY502924, AY278491, and AY502927) were aligned by MEGA7 software.[22] SARS-CoV-2 specific regions for LAMP primer design are manually selected: two regions from *Nsp3* (3055 – 3591, 6172 – 7273), two regions from *Spike* (S) (21540 – 22549, 22890 – 23779), and one region from *Orf8* (27824 – 28396). Whole *Nucleocapsid* (N) gene region is also included as *N* is usual target of molecular diagnosis due to abundance of its mRNA.[23] Two to five basic LAMP primer sets are designed and selected with PrimerExplorer V5 (http://primerexplorer.jp/lampv5e/index.html) for each target region. Loop primers are designed by PrimerExplorer V5 or manually selected. 16 LAMP primer sets with proper both of Loop Forward (LF) primer and Loop Backward (LB) primer are selected and subjected to further screening.

### LAMP Primer Screening

First round of screening was done using WarmStart Colorimetric LAMP 2X Master Mix with cDNA of which corresponding RNA concentration is 8.3 × 10^4^ copies/reaction. Nine out of 16 primer sets were selected by threshold time from the results of 40 minutes incubation at 65°C and subjected to further screening.

Second round of screening was done using *Bst*3.0. The same amount of cDNA as the first round of screening is used. Two primer sets that showed relatively early non-specific amplification are discarded and remaining seven primer sets are subjected to further screening.

Third and fourth rounds of screening were done by checking sensitivity to dilutions of cDNA and RNA, respectively. Five out of seven primer sets showed specific amplification for at least one replicate of duplicate with cDNA concentration corresponding to 1.7 × 10^1^ copies of input RNA (Supplementary Figure 1). Next, we evaluated sensitivity of the five primer sets to RNA dilutions in RT-LAMP using *Bst*3.0 and SuperScript IV Reverse Transcriptase. Two primer sets, both targeting *Nsp3*, showed best sensitivity that showed specific amplification at 10^−6^ dilution of RNA (Figure 1). We also added two primer sets, one targeting *S* and the other targeting *N*, for reaction optimization experiments as they showed fast threshold time for cDNA and to keep ranges of target genes. Primer sequences are represented in Table 1.

**Table 1.**
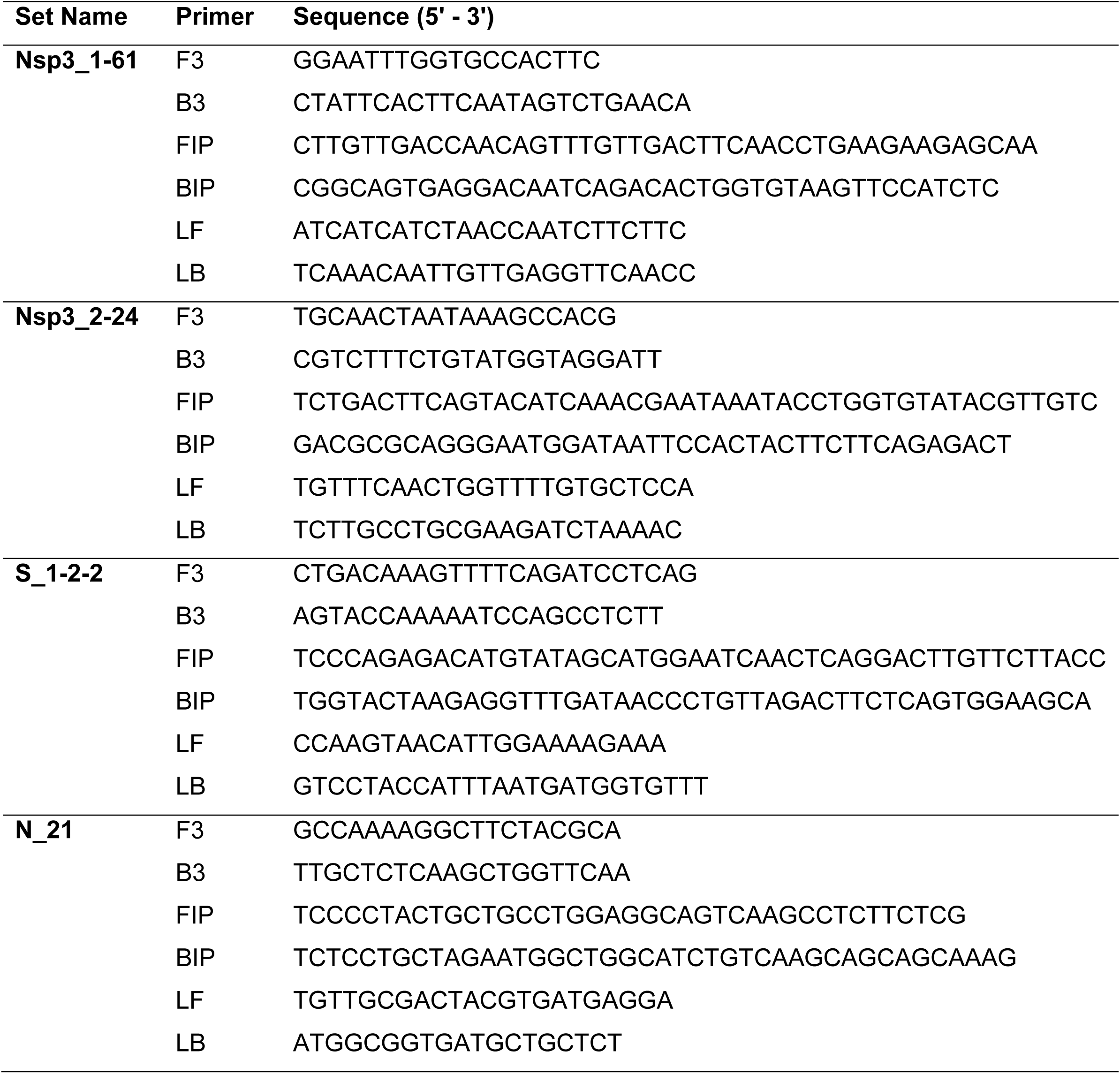
LAMP primers of which assays optimized and LoD evaluated.

**Figure 1.**
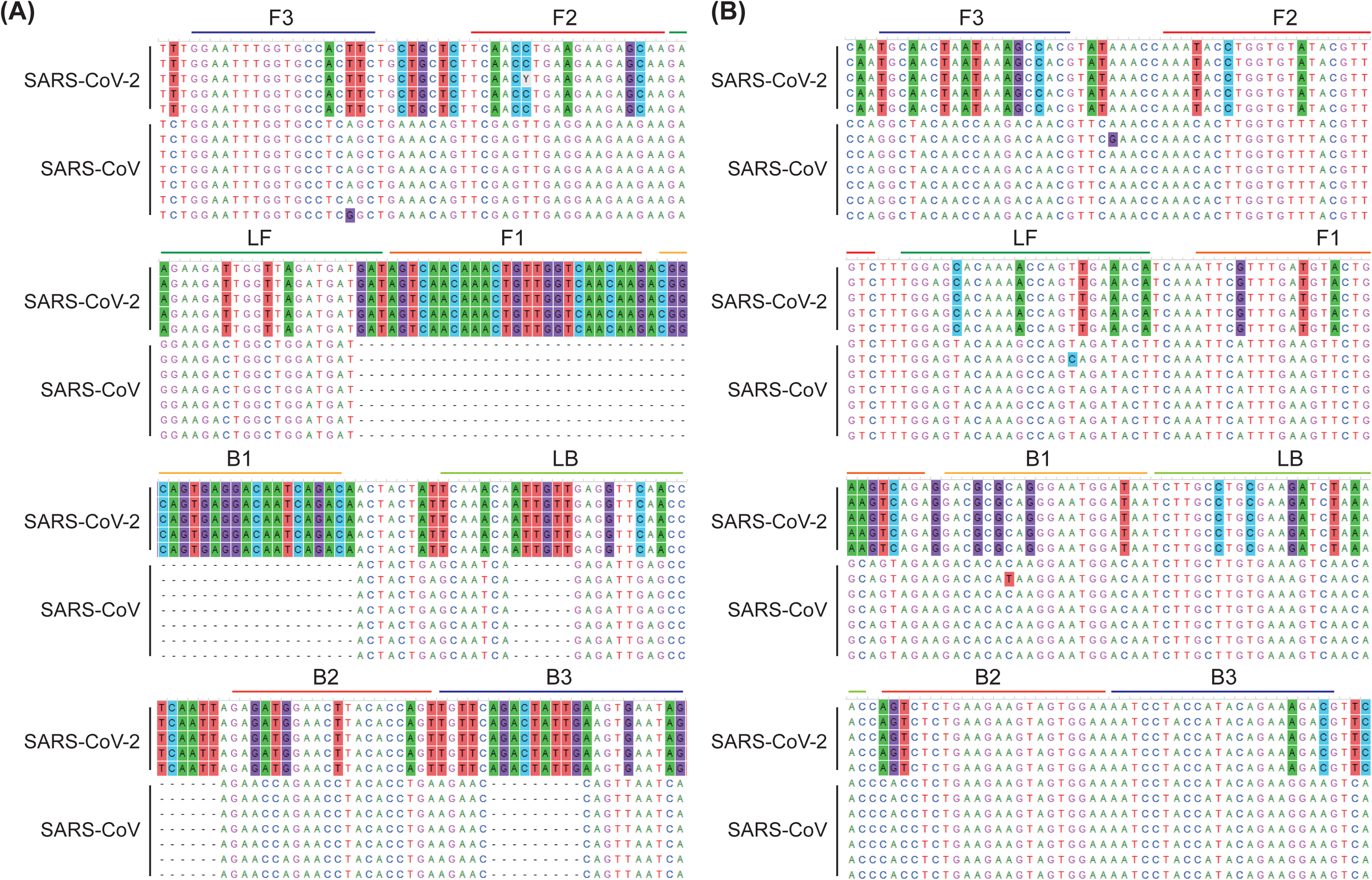
LAMP primer positions on aligned sequences of SARS-CoV-2 and SARS-CoV Primer binding sites of (A) “Nsp3_1-61” and (B) “Nsp3_2-24” LAMP primer sets are depicted on aligned sequences of five SARS-CoV-2 (from up, MN908947, MN938384, MN988713, MN985325, MN975262) and seven SARS-CoV (from up, NC_004718, AY613947, AY313906, AY559094, AY502924, AY278491, AY502927). Conserved sites are toggled at 50% level by MEGA7 software so that SARS-CoV-2 specific residues versus SARS-CoV are background colored.

### LAMP Reaction Optimization

To optimize RT-LAMP reaction with *Bst*3.0, we first evaluated optimal concentration of dNTP and Mg^2+^ for each primer set. Three dNTP-Mg^2+^ concentration combinations are tested at 69°C: dNTP 1.4 mM and Mg^2+^ 8 mM, dNTP 1.4 mM and Mg^2+^ 6 mM, dNTP 1 mM and Mg^2+^ 6 mM. The copy number of RNA template was 1,000 copies per reaction. Best concentration combination was selected for each primer set by whether both of duplicate show specific amplification and by threshold time. Average threshold time (Tt_av) and corresponding dNTP and Mg^2+^ concentrations are as follows: dNTP 1 mM and Mg^2+^ 6 mM for “Nsp3_1-61” (Tt_av = 11.93), dNTP 1.4 mM and Mg^2+^ 8 mM for “Nsp3_2-24” (Tt_av = 7.74), dNTP 1.4mM and Mg^2+^ 6mM for “S_1-2-2” and “N_21” (Tt_av = 11.08 and 5.50, respectively).

Next, we checked if amplification is improved at lower temperature (65°C) because optimal reaction temperature of SuperScript IV Reverse Transcriptase is 50-55°C and manufacturer recommend 65-72°C for optimal performance of *Bst*3.0. Notably, “Nsp3_1-61” and “S_1-2-2” primer sets show improved threshold time (Tt_av = 8.92 and 8.49, respectively).

### Assessing Limit of Detection and Cross-Reactivity

The Limit of Detection (LoD) of optimized RT-LAMP assays were assessed through 5 to 1000 RNA copies/reaction in triplicate. Among four primer sets subjected to reaction optimization, “S_1-2-2” and “N_21” sets showed relatively poor sensitivity. For “Nsp3_1-61” and “Nsp3_2-24” primer sets, we additionally evaluated LoD with 2-fold SuperScript IV Reverse Transcriptase (20U/reaction). LoD and Tt_av were improved by increasing reverse transcriptase amount. (Table 2, Supplementary Figure 2) As a result, both “Nsp3_1-61” and “Nsp3_2-24” RT-LAMP assays could detect as low RNA concentration as 100 copies per reaction.

**Table 2.** LoD of RT-LAMP assays targeting SARS-CoV-2.

| RNA<br>copies | SSIV, 10U/rxn |  |  |  | SSIV, 20U/rxn |  |  |  |
| --- | --- | --- | --- | --- | --- | --- | --- | --- |
|  | Nsp3_1-61 |  | Nsp3_2-24 |  | Nsp3_1-61 |  | Nsp3_2-24 |  |
|  | Tt_av | RPR* | Tt_av | RPR* | Tt_av | RPR* | Tt_av | RPR* |
| **1000** | 13.11 | 3/3 | 9.31 | 3/3 | 10.7 | 3/3 | 7.53 | 3/3 |
| **500** | 13.47 | 3/3 | 8.16 | 2/3 | 10.63 | 3/3 | 7.59 | 3/3 |
| **200** | 13.65 | 2/3 | 11.19 | 2/3 | 11.34 | 2/3 | 7.78 | 3/3 |
| **100** | 14.62 | 3/3 | - | - | 12.26 | 2/3 | 8.47 | 2/3 |
| **50** | - | - | - | - | - | - | - | - |
| **20** | 13.88 | 1/3 | - | - | - | - | - | - |
| **5** | - | - | - | - | - | - | - | - |
| **NTC** | - | - | - | - | - | - | - | - |

\* RPR : Ratio of Positive Replicates

The cross-reactivity of two SARS-CoV-2 RT-LAMP assays targeting *Nsp3* to other human Coronaviruses was not found for hCoV-229E, hCoV-OC43 and MERS-CoV (Figure 2).

**Figure 2.**
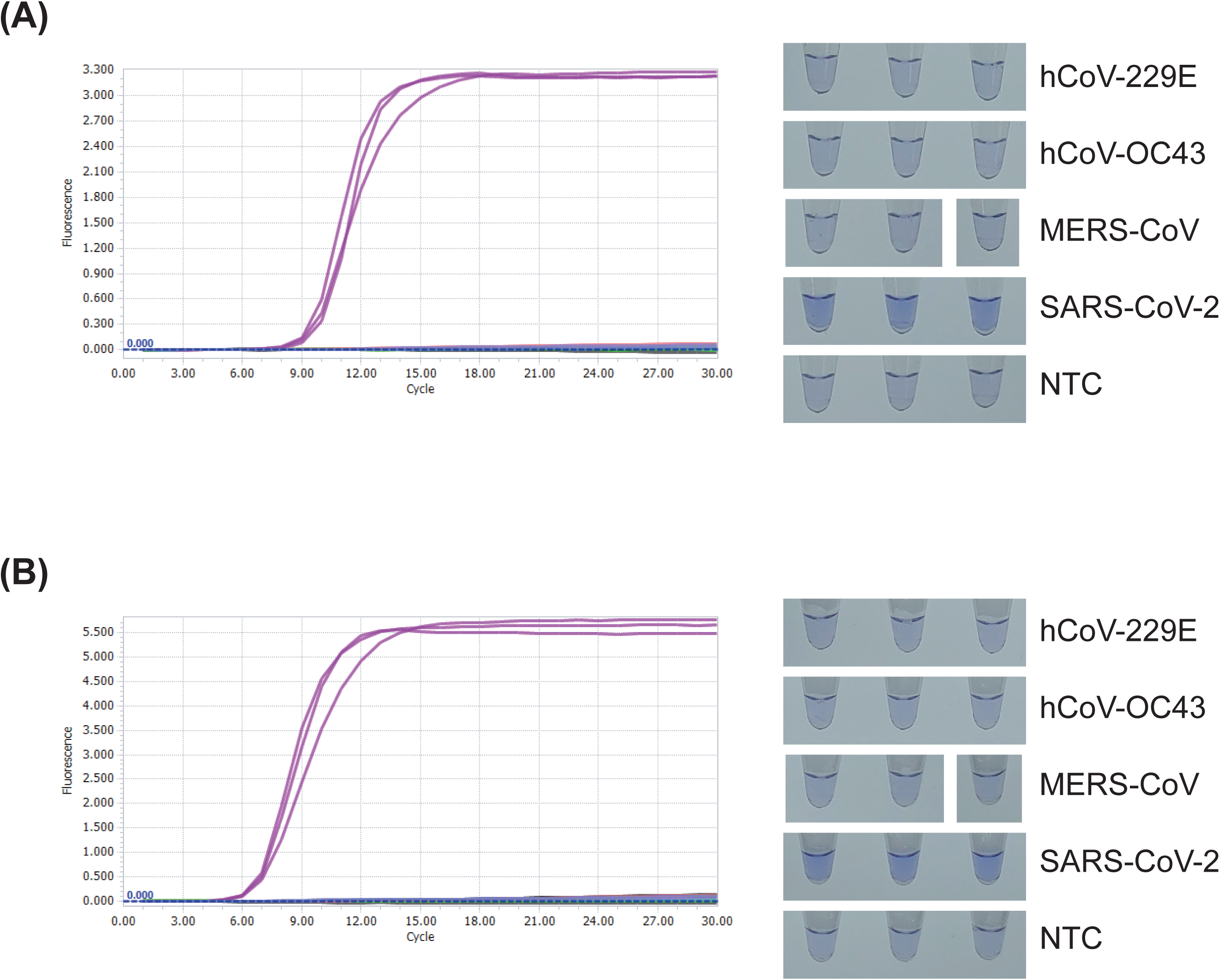
Cross-reactivity to other Coronaviruses tested for RT-LAMP assay targeting SARS-CoV-2 Real-time amplification fluorescence signal and end-point LCV colorimetric detection results of cross-reactivity test for (A) “Nsp3_1-61” and (B) “Nsp3_2-24” LAMP primer sets. Reactions are performed with 20U/reaction of reverse transcriptase and optimized temperature, dNTP concentration and Mg^2+^ concentration for each primer set. RNA copy number of each viral RNAs are as follow: hCoV-229E – 1.6 × 10^6^, hCoV-OC43 – 1.6× 10^6^, MERS-CoV – 4.5× 10^6^, SARS-CoV-2 – 2.5 × 10^3^.

## Discussions and Conclusion

In this study, we designed and screened LAMP primer sets targeting SARS-CoV-2. Reaction optimization was also performed for selected primer sets. In summary, we designed and evaluated a pair of RT-LAMP assays for detection of SARS-CoV-2 with limit of detection of 100 copies per reaction. Our RT-LAMP assays showed specificity to SARS-CoV-2 versus alphacoronavirus (hCoV-229E), betacoronavirus (hCoV-OC43) and MERS-CoV. Although we could not test specificity to SARS-CoV because proper sample was not in our hand, specificity of the RT-LAMP assays are easily expectable from the mismatching bases in primer binding sites (Figure 1). Especially, both of F1/B1 sites of “Nsp3_1-61” primer set is in SARS-CoV-2 specific region of which aligning SARS-CoV sequence is not exist.

About the sensitivity of LAMP assays, note that average threshold time is not well correlated with limit of detection. In fact, average threshold time of “S_1-2-2” and “N_21” primer sets for 1000 copies were faster than that of “Nsp3_1-61”: 7.33 (RPR: 2/3) and 5.06 (RPR: 1/3) minutes, respectively. This dis-relation is previously reported [24].One peculiar observation is that both “S_1-2-2” and “N_21” showed good sensitivity to cDNA template (Supplementary Figure 1) unlike to RNA. Observed difference of sensitivity to RNA and cDNA seems significant even we account slight amplification that might happened during reverse transcription. The reason may include stochastic nature of forming proper LAMP amplification intermediate - “dumbbell” structure [24], higher stability of DNA:RNA double strand than DNA:DNA double strand and more.

The limit of detection of RT-LAMP assays suggested in this study, 100 copies per reaction, may not enough for sensitive screening of suspicious patients. This relatively high limit of detection would be from the target sequences used for primer design that are selected by specificity versus SARS-CoV. Indeed, target GC percentage and Tm of primers had to be adjusted to get enough number of starting sets for primer screening during LAMP primer design to give less-optimal primer sets. Therefore, there might be better target sequences for LAMP assay of SARS-CoV-2 in the view point of sensitivity. However, expected high specificity of RT-LAMP assays suggested in this study would be a good feature for a confirmatory test. In addition, considering high viral load of SARS-CoV-2 at early stage after symptom onset [25], suggested RT-LAMP assays may still be usable for screening tests.

In conclusion, we developed highly specific RT-LAMP assays for detection of SARS-CoV-2. The results of these RT-LAMP assays can be detected within 30 minutes after amplification reaction begin. In addition, we provided optimized reaction conditions to which LCV colorimetric detection method is applied that can be used for point-of-care tests.

## Acknowledgements

This work was supported by the National Research Council of Science and Technology (NST) grant by the Ministry of Science and ICT (Grant No. CRC-16-01-KRICT).

---

**Supplementary Figure 1.**
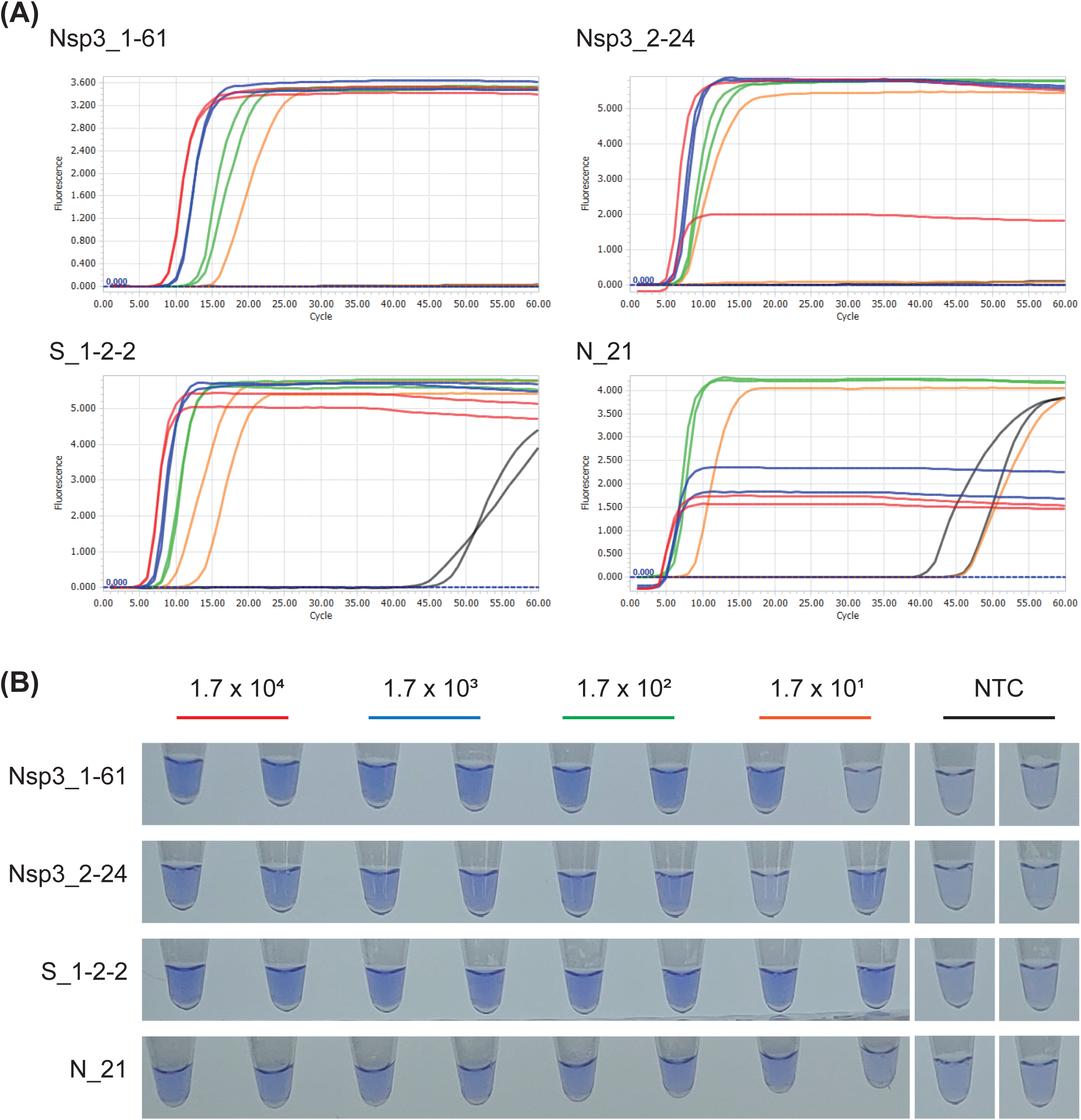
Sensitivity of SARS-CoV-2 LAMP assays to cDNA dilutions. (A) Real-time amplification fluorescence signal and (B) end-point LCV colorimetric detection results of four SARS-CoV-2 LAMP primer sets that were subjected to reaction optimization. Color of amplification signal curve and LCV detection label for each cDNA dilution are matched. Designated copy number is corresponding reverse transcription input RNA copy number. Some low fluorescence of amplification signal plateaus are due to baseline correction process of LightCycler 96 software.

**Supplementary Figure 2.**
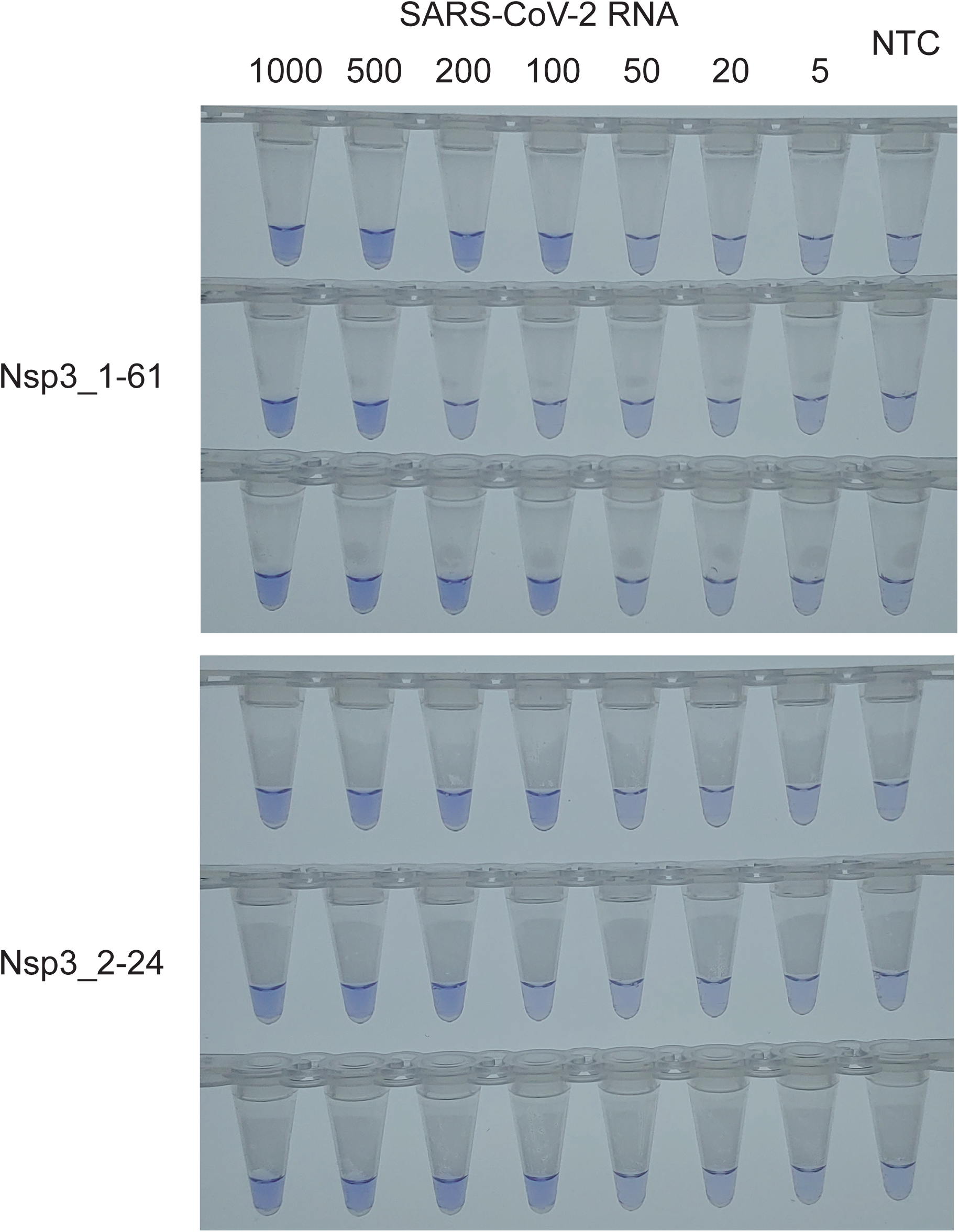
Colorimetric detection of SARS-CoV-2 RT-LAMP LoD tests for “Nsp3_1-61” and “Nsp3_2-24”. LCV colorimetric detection results of LoD tests for “Nsp3_1-16” and “Nsp3_2-24”. 20U/reaction of reverse transcriptase were used. See main text for detailed reaction condition.

